# Evolutionary puzzle: discussing the evolution of sentience in Metazoa through a phylogenetic perspective

**DOI:** 10.1101/2021.05.21.445182

**Authors:** Michaella P. Andrade, Charles Morphy D. Santos

## Abstract

Sentience is the capacity of organisms to feel and experience through subjective states. During the last years, several investigations have indicated that response mechanisms to harmful stimuli can be highly conserved among the Metazoa. However, there is a research bias towards vertebrates in the available studies. Here we discuss the evolution of the nervous and sensory system, pain and nociception in animals through a phylogenetic perspective testing the hypothesis of common ancestry of sentience. Our results indicate that characteristics related to sentience - morphological and molecular and behavioural -, were already present in the common ancestors of Metazoa, Eumetazoa and Bilateria. Our phylogenetic hypotheses positioned Porifera as the sister-group to all the other Metazoa, corroborating the hypothesis of a single origin of the nervous system. Our results also depict Urbilateria as the ancestor of the metazoan toolkit related to the sentience. These scenarios suggest that some attributes of the sensory system may have appeared even before the emergence of the nervous system, through possible cooptations of sensory modules of the first Metazoa.

## 1. Introduction

Sensory capabilities are biological phenomena of great ecological and evolutionary relevance, playing important roles in evolutionary radiation, coevolution and generation of biodiversity (Muller et al. 2018). A better understanding of the sensory capacities of animals could help us to understand the evolution of behavioural plasticity of responses to various stimuli in unstable environments, and to comprehend the origin and diversification of sentience in nature (Godfrey-Smith 2017; Feinberg and Mallat 2019).

Sentience is the capacity of organisms to feel, sense and experience positively or negatively through subjective states (Proctor 2012; Broom 2016; Bronfman et al. 2016; Godfrey-Smith 2016; Feinberg and Mallat 2019; Birch 2020). Every animal with some capacity to feel, recognize and be affected by stimuli has the potential to be sentient (Bronfman et al. 2016; Godfrey-Smith 2016). Beyond humans, studies have suggested the presence of sentience in other amniotes (such as mammals and birds), other vertebrates and some protostomes (as cephalopods) (Macphail 1998; Edelman and Tononi 2000; Seth et al. 2005; Cabanac and Parent 2009; Ginsburg and Jablonka 2010; Griffin 2013; Sovik and Perry 2016). Some authors even propose the widespread presence of sentience in the whole tree of life (Margulis 2001; Baluska and Reber 2009; Pereira 2016).

One major challenge is to define criteria to establish the actual presence of sentience (Harnad 2016). Since non-human animals do not use human language, what these animals are feeling can only be inferred through indirect criteria (e.g. physiological, evolutionary, behavioural) - this is the well-known “other minds problem” (Harnad 2016; Sovik and Perry 2016; Gutfreund 2018; Birth 2019; Feinberg and Mallat 2019; Godfrey-Smith 2019). To determine the real frontiers of the sentience and its constituent blocks remains an inconclusive question (Smith et al. 2009; Proctor 2013; Teles 2016; Veit and Huebner 2020).

Recently, two hypotheses were suggested to explain the evolution of sentience: convergences in several clades of Metazoa (Feinberg and Mallat 2016, 2019, 2020) or shared ancestry (Sovik and Perry 2016). In the first hypothesis, the sentience must have independently evolved in Arthropoda, Cephalopoda and Vertebrata, which are the metazoan clades that presenting characteristics that allow the existence of subjective states (e.g. sensory organs, distinct types of neurons and more brain regions, effective memories and operant learning). According to this scenario, the centralization of the nervous system occurred independently among such lineages from a common Bilateria ancestor with a net nervous system (Feinberg and Mallat, 2016).

In the hypothesis of homology of sentience among different taxons of Metazoa, the last common ancestor presenting some degree of sentience possibly have had a centralized nervous system allowing the integration of sensory stimuli and motor outputs. In this sense, this phenomenon could not have existed outside Bilateria (Sovik and Perry 2016), which would bring the origin of sentience to the beginning of the Phanerozoic, ca. 540 million years ago (Sovik and Perry 2016). Evidence that supports such a hypothesis are molecular and morphological similarities (i.e. deep homology) among the central nervous system of Arthropoda and the basal ganglia of vertebrates (Strausfeld and Hirth 2013). In this scenario, Urbilateria would already have a nervous system with ganglionic clusters. This putative ancestor is known as the Last Universal Common Sentient Ancestor (LUCSA).

Although great advances have been achieved in recent decades, the aforementioned studies do not exhaust the subject. In the present paper, we discuss scenarios for the evolution of sentience in animals based on a phylogenetic framework. We use different sets of characters - morphological, molecular and behavioural - and reciprocal illumination techniques to test the hypothesis of common ancestry of sentience since the origin of the first organisms characterized by the presence of bilateral symmetry and its associated attributes. Regarding sentience, there seems to have been few evolutionary paths towards the plethora of particular solutions at all levels of the tree of life. The main aim here is to discuss how narrowly circumscribed life is when it comes to the evolution of the animal capacities to feel, sense and experience subjective states.

## 2. Material and Methods

### 2.1. Taxon sampling

Our phylogenetic analyses based on morphological data (Table S1 and S2) include 26 supra-specific clades of Metazoa and related groups (Actinopterygii, Annelida, Amphibia, Birds, Bivalvia, Chelicerata, Cephalochordata, Cephalopoda, Choanoflagellata, Chondrichthyes, Cnidaria, Crocodilia, Crustacea, Ctenophora, Echinodermata, Gastropoda, Hexapoda, Lepidosauria, Mammalia, Myriapoda, Nematoda, Platyhelminthes, Polyplacophora, Porifera, Testudinata and Urochordata). The analyses based on molecular data include 24 of the supra-specific terminals used in the morphological analyses with the exclusion of Myriapoda and Polyplacophora (see details in Table S3).

### 2.2. Morphological, molecular and behavioural data

The morphological characters were coded by attributes identified in the literature resulting in a matrix with 122 characters (Table S1). In the molecular analyses, we selected nucleotide sequences of eleven genes, nine of them directly involved in nociception and pain mechanisms, and two markers recognized as homologous among animals: DegEnac, NomP, Prdm12, Pkd2, Piezo, TRPV1, TRPA1, TRPC3, TRPM8, Chordin/Sog, and 18S (Table S3). All the sequences were downloaded from the GenBank Platform (NCBI) constituting a database with 23.301 molecular bases.

We compiled a survey of behavioural data related to sentience, and ploting the presence of two types of associative learning, classical and operant conditioning, and the ability to respond to three distinct stimuli types (mechanical, thermal and chemical) in Metazoan taxa at our tree obtained phylogenetic analysis based on morphological characters (Table S4, Table S5 and Table S6).

### 2.3. Phylogenetic analyses

The morphological data matrix was constructed in Winclada (Nixon 2002) (Table S1) and the parsimony analysis was performed through TNT software (Goloboff and Catalano 2016). Heuristic searches were performed with the TBR algorithm, with 500 replicas and 10 trees saved each. We used ACCTRAN optimization (Agnarsson and Miller 2008) to visualize the characters and their states in the most parsimonious final tree. Consistency (CI) and retention indices (RI) (Farris 1989) of the tree and the individualized characters were used as tools for understanding the results obtained (Figure S1). Bremer support measures (decay rates) (Bremer 1994) were also presented to identify the stability of our resultant clades. Outgroup comparison was used to polarize the characters (Nixon and Carpenter 1993; Bryant and Wagner 2001); here, the chosen outgroup was the opisthokont Choanoflagellata, used on a regular basis in phylogenetic analysis of Metazoa (e.g. Sorensen et al. 2000; Edgecombe et al. 2011; Dunn et al. 2014).

The molecular sequences went through multiple alignments with ClustalW in Bioedit (Hall 1999). After edition, each molecular sequence was concatenated using the Merge Matrices Tool in Winclada software. The phylogenetic relationships were obtained through parsimony, maximum likelihood and Bayesian inference. Parsimony analyses were performed in TNT (Goloboff et al. 2016), with 1000 replicas and 10 trees saved for each one, and the strict and majority consensus (50%) were computed for the equally most parsimonious trees found. For Maximum Likelihood inference (Felsenstein 1981), the sequences were transferred to CIPRES Science Gateway platform (Miller et al. 2010) in RAxML v8.1 program (Stamatakis 2014). Only one partition was selected by PartitionFinder2 under the GTR + G model, with Choanoflagellata as the outgroup. The support measure Rapid Bootstraps (Stamatakis et al. 2008), with 1000 replicates, was used to estimate the reliability of each node of the final topology. In the Bayesian Inference (Rannala and Yang 1996; Huelsenbeck et al. 2001), we partitioned our data in PartitionFinder2 (Lanfear et al. 2012) in CIPRES (Miller et al. 2010) to obtain the best evolutionary replacement models of each subset with the greedy algorithm (Lanfear et al. 2016). Bayesian searches were conducted in Mr. Bayes 3.2 (Ronquist et al. 2012) with parameters 2 and 4 races and 10 million generations, discarding the first 25% trees as burn-in (Lemey et al. 2009).

## 3. Results

### 3.1. Phylogenetic analysis based on morphological data

The phylogenetic analysis of the morphological data matrix resulted in a single most parsimonious tree, with 152 steps, CI=0.88, and RI=0.95. Eumetazoa, Bilateria, Deuterostome, Protostome, Ecdysozoa, Spiralia, Chordata, Vertebrata and Amniota were recovered as monophyletic groups, being the highest values of the Bremer support measure found in Metazoa, Eumetazoa, Vertebrata and Ecdysozoa (Figure 1).

**Figure 1.**
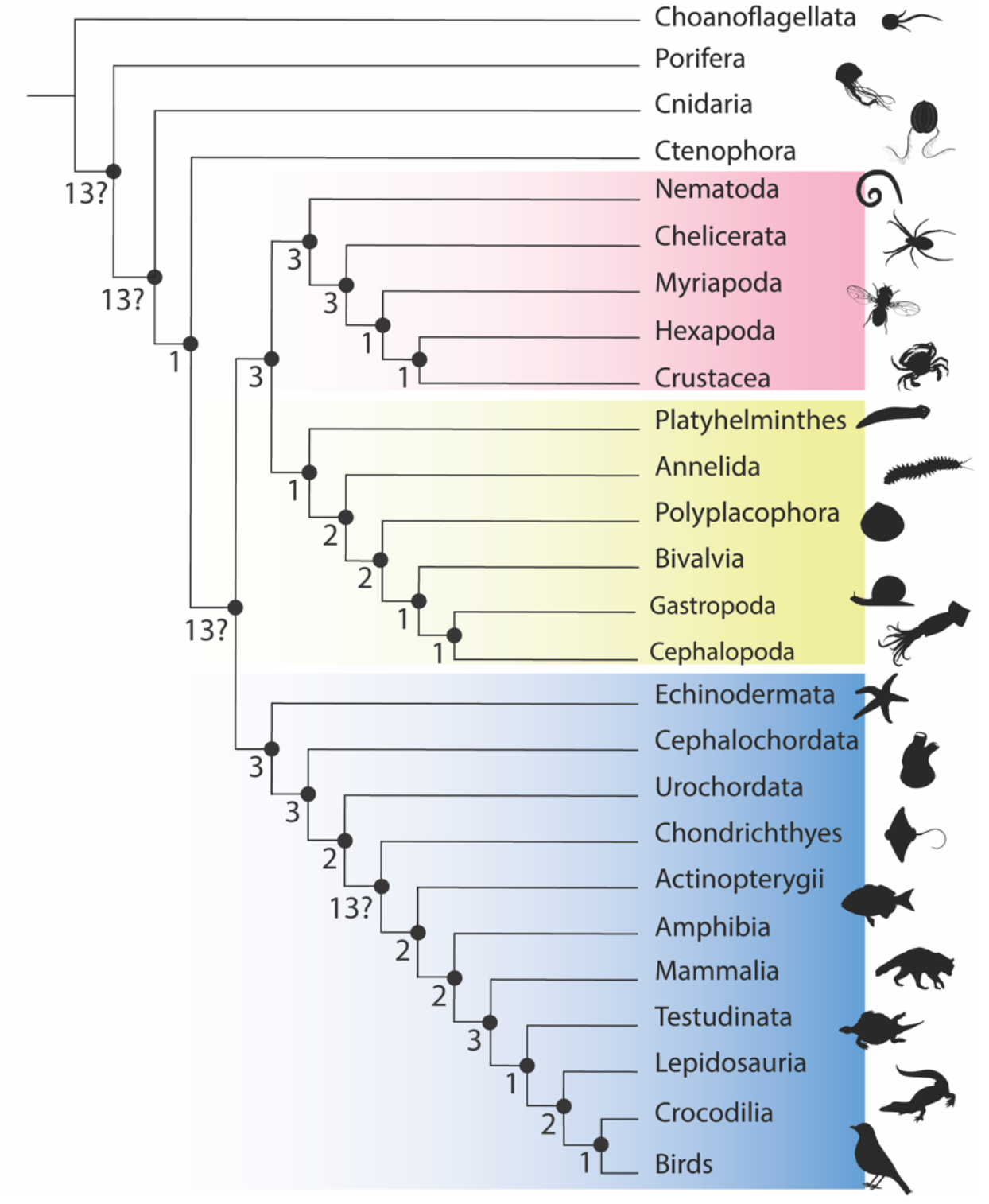
Phylogenetic relationships of the main groups of Metazoa obtained by the parsimony analysis of 122 morphological characters. Bremer support measure pointed on the nodes of each grouping. The blue boxes indicate Deuterostomia, the yellow boxes Spiralia and pink boxes Ecdysozoa. Silhouette of animals were downloaded from *http://phylopic.org*.

According to our results, the common ancestor of Metazoa presented multicellularity, collagen, intermediate filaments, microfilaments, spermatogenesis, actin, myosin, primary sensory cell and presence of immobile cilia in the sensory cell, besides neurotransmitters such as Glutamate, GABA and Histamine. The Eumetazoa was also defined through undisputed synapomorphies: presence of true tissues, basal membrane, specialized gonads, epithelia with polarized cells joined by cellular junctions, Hox genes, nervous system, endogenous opioid receptors, statocyst, microvilliates present in sensory cells, support cells in mechanoreceptors, and the neurotransmitters Glycine, Dopamine, Norepinephrine, Epinephrine and Serotonin. Embryonic leaflets and symmetry patterns are also widespread in Eumetazoa, with reversions.

Based on our phylogenetic hypothesis, in bauplan of Bilateria were present an excretory system, commissure between the nervous cords, nociceptors, amiellinic nerve fibers, glial cells and cerebral ganglion. Characters such as the presence of celloma, circulatory system, bilateral symmetry, an organized nervous system and nerve cords, with different degrees of variation, also date from the origin of Bilateria.

The two clades of Bilateria - Protostomia and Deuterostomia - were recovered as monophyletic groups in our phylogenetic hypothesis. Protostomes are mainly characterized by the presence of cleavage in the superficial state, some sort of hemocele, neurotransmitter octopamine and subepidermal ventral nervous system. Deuterostomes was sustained by the characters formation of the celloma, blastopore destiny and mouth originating from the gastrodermis, pharynx with ciliated gill slits, organization of the nervous system, originated from invaginations, in the dorsal region.

### 3.2. Phylogenetic analysis based on molecular data

Our results were obtained through parsimony analysis, maximum likelihood and Bayesian inference. Despite the method, phylogenetic hypotheses based on molecular evidence were less resolved than hypothesis based on morphological data and openly incongruent with the literature. In fact, our main aim here was to identify molecular sequences or toolkits that may have originated long before the emergence of sentience, possibly related to the first processes of nociception and pain mechanisms in ancient animals. In this sense, the molecular markers were sufficient to support the main clades within Metazoa. This strongly suggests that the common ancestors of groups such as Eumetazoa and Bilateria were at least partially equipped with the general molecular attributes that would be diversified in processes related to animal sentience.

The parsimony analysis of the molecular sequences gathered in our study resulted in 10 equally most parsimonious trees. Both strict and majority consensus trees indicated Porifera as the sister-group to all of the remaining Metazoa, suggesting that Eumetazoa is a monophyletic group. As well as our morphologically based topology, the analyses recovered traditional groups such as Bilateria and Deuterostomia (including here Chordata, Vertebrata and Amniota). The positioning of certain terminals - such as Hexapoda and sister-group to Nematoda, and Crocodilia as sister-group to Testudinata - is not congruent with the literature (e.g., Chiari et al. 2012; Dunn et al. 2014; Giribet and Edgecombe, 2020). Still, Crustacea, Platyhelminthes and Urochordata are *incertae sedis* according to both consensus.

As expected, the strict consensus (Fig 2, A) is poorly resolved. Although Ecdysozoa (pink box) has been partially recovered, the positioning of Crustacea is enigmatic. Chordata was recovered as monophyletic similar to the result found in the analysis based on morphological data. In the majority consensus (Fig 2, B), Bilateria is monophyletic, but its internal relationships are not completely resolved. Spiralia (yellow box) was partially recovered. Deuterostomia is also a natural group, with Echinodermata as the sister group to the other deuterostomes (although Urochordata remained outside this group). Part of the terminals traditionally belonging to Protostomia are not definitively positioned, forming a polytomy with other terminals in both strict (Fig. 2, A) and majority consensus (Fig 2, B).

**Fig. 2.**
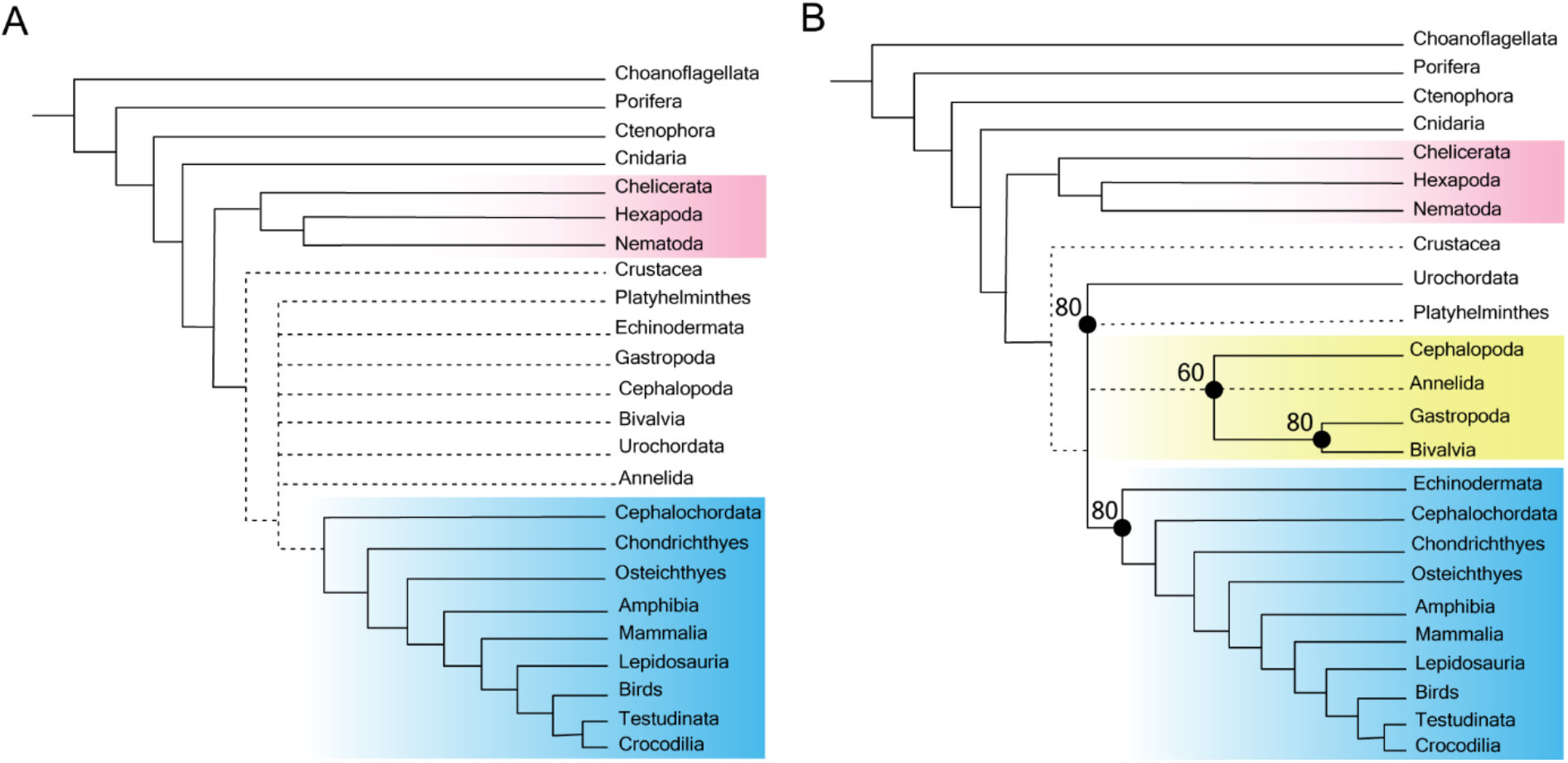
Phylogenetic hypotheses based on parsimony analysis of the molecular data. (A) the strict consensus, and (B) majority consensus of 10 equally most parsimonious trees.

The main clades of Metazoa were recovered in our Maximum Likelihood analysis - Eumetazoa, Bilateria, Protostomia, Spiralia, Deuterostomia, Chordata and Vertebrata (Fig 3, C). Despite Chordata, the internal relationships of these main clades were mostly poorly resolved or incongruent to the results obtained with the morphological analysis, with low Bootstrap values. A similar scenario resulted from the Bayesian Inference (Fig 3, D). Here, we obtained the highest subsequent probabilities for the traditional clades within Metazoa, with Cnidaria as sister-group to Bilateria.

**Figure 3.**
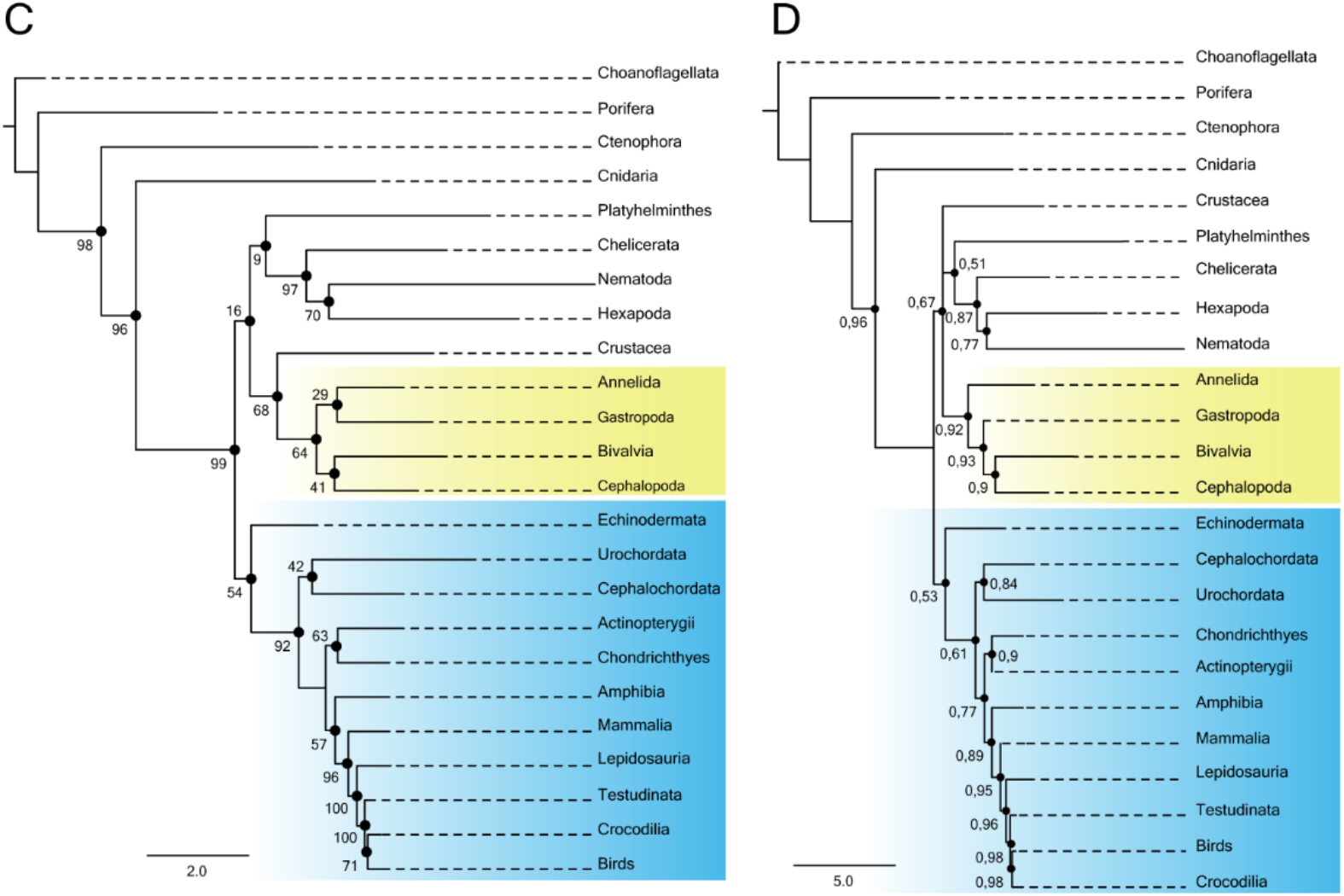
Phylogenetic hypotheses based on (C) Maximum Likelihood with rapid Bootstraps values pointed at the node and (D) Bayesian Inference with probabilities higher than 50 pointed at the nodes.

### 3.3. Distribution of behaviours in the phylogeny of Metazoa

Plotting behavioural characters related to sentience - presence of classical and operant conditioning, and the ability to respond to mechanical, thermal and chemical stimuli - on our morphological phylogeny resulted in the topology presented in Figure 4. Classic conditioning is widely present among animals, while the operant conditioning was reported for some groups of Mollusca (Gastropoda and Cephalopoda) and Arthropoda (Hexapoda and Crustacea); there are insufficient data for Ctenophora, Myriapoda, Polyplacophora, Bivalvia, Cephalochordata, Urochordata and Chondrichthyes. Operant conditioning was recovered with restricted distribution in Hexapoda, Crustacea, Gastropoda, Cephalopoda and Vertebrata. Behavioural responses to harmful mechanical and chemical stimuli are widespread in Metazoa (even in organisms lacking nervous systems such as Porifera). Behavioural responses to harmful thermal stimuli appear in Cnidaria and some Bilateria, although data for Ecdysozoa, Spiralia and ancient lineages are missing.

**Figure 4.**
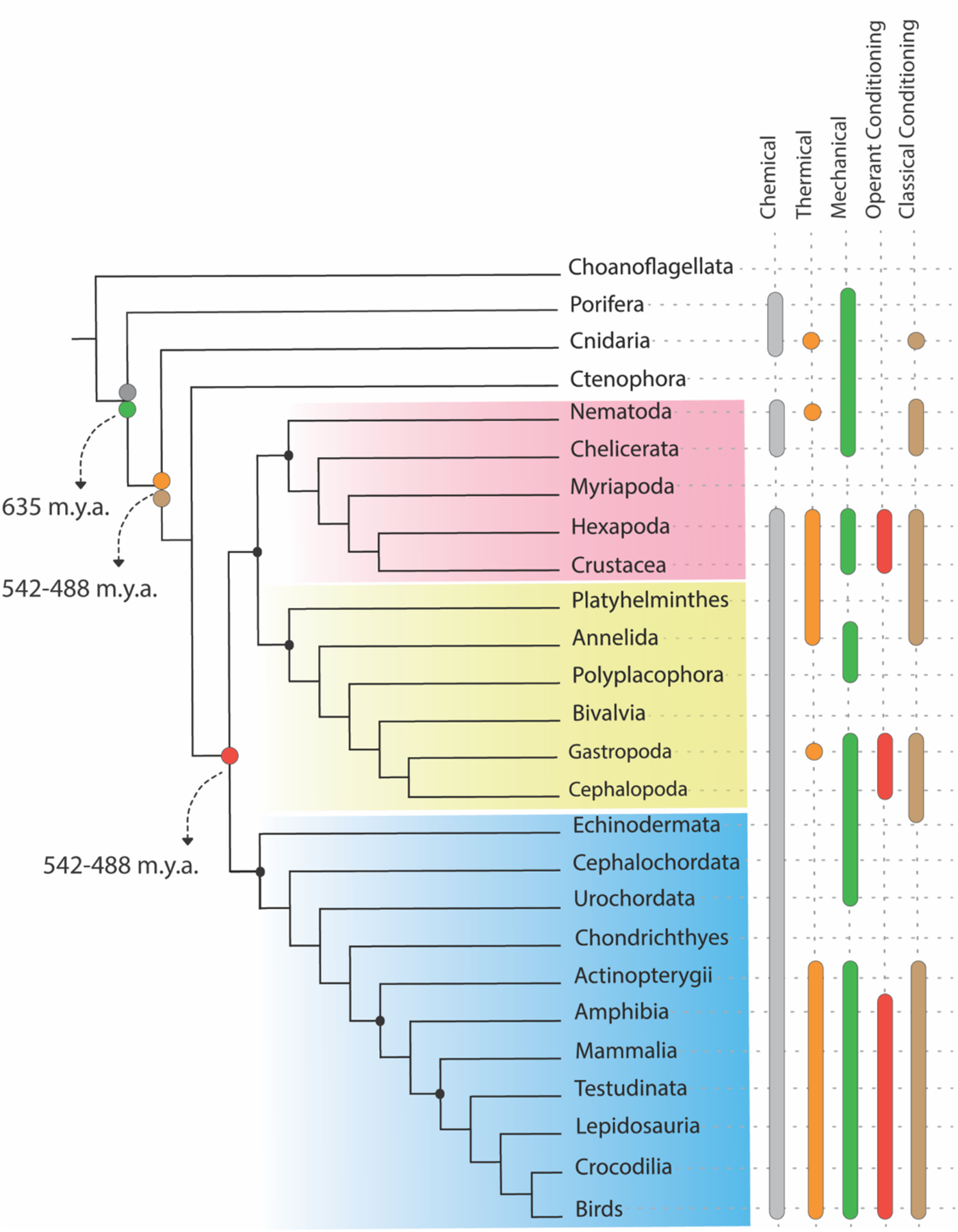
Hypothesis of phylogeny of Metazoa with the optimization of the presence of five behaviours with time of divergence of these lineages according to Giribet and Edgecombe (2020).

## 4. Discussion

The phylogenetic continuity of animal sentience is a controversial topic (Ginsburg and Jablonka 2010; Godfrey-Smith 2017; Feinberg and Mallat 2019). However, evolutionary biology has taught us that all animals species share several degrees of their evolutionary history, and so we may wonder why cognitive abilities could not follow, at some level, the same path (De Wall 2019). Assuming that sentience is related to different neuronal cell modifications and associated processes (Ginsburg and Jablonka 2010; Sovik and Perry 2016; Godfrey-Smith 2017; Feinberg and Mallat 2019), our results support the LUCSA hypothesis. According to such hypothesis, sentience is widespread among true bilateral animals strains with true bilateral symmetry (Figure 5). However, even before the origin of the last universal common sentient ancestor (and the emergence of nerve cells and centralized nervous systems), plesiomorphic conditions related to sentience were already present.

**Fig. 5.**
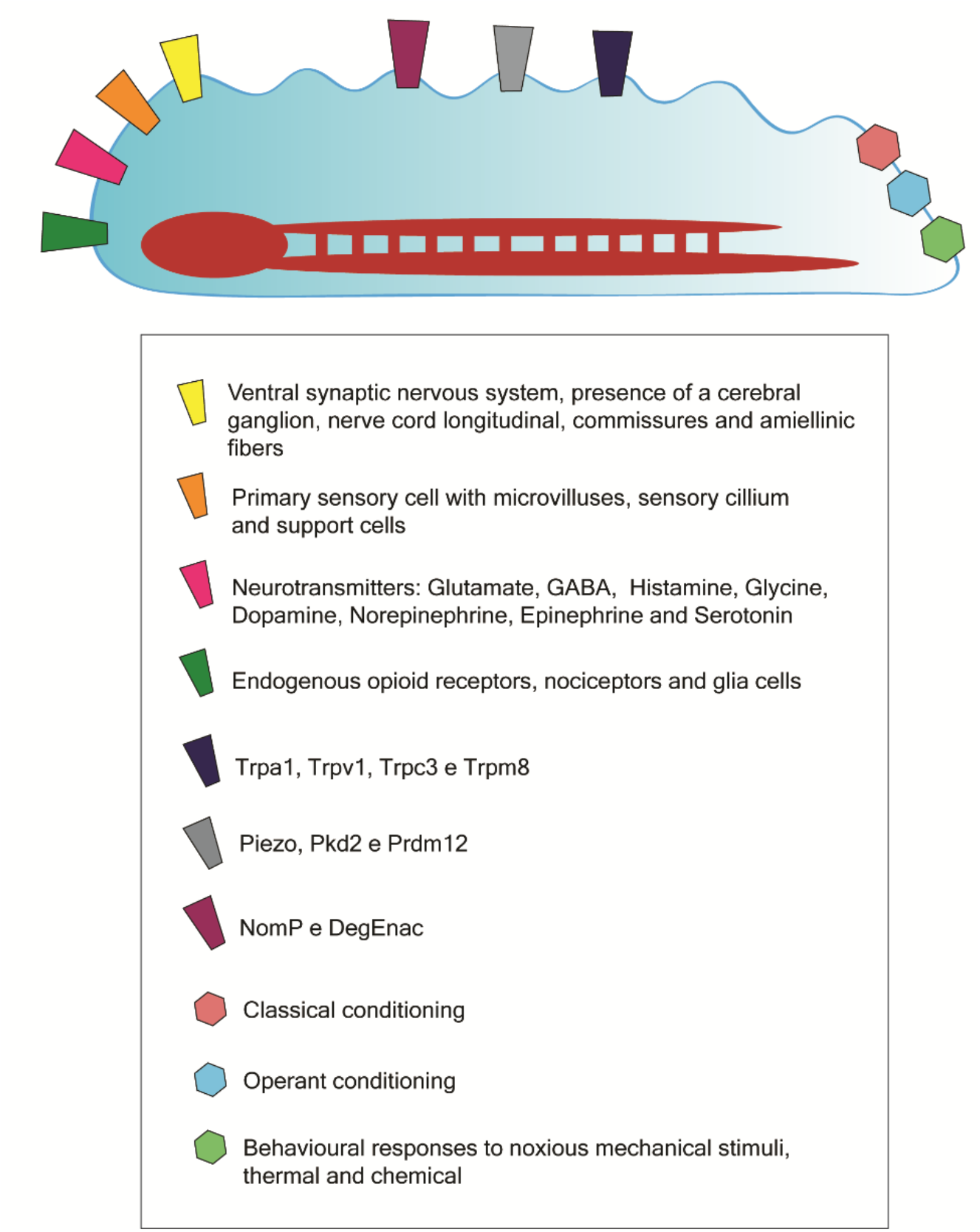
Schematic reconstruction of Urbilateria, the hypothetical ancestor of all of the bilaterian animals, with the morphological, molecular and behavioural attributes discussed here.

### 4.1. Characters supporting the LUCSA hypothesis

According to our results, neurotransmitters such as GABA, Glutamate and Histamine (Karhunen et al. 1993; Liebeskind et al. 2017; Byrne 2019) are omnipresent in Metazoa, including Porifera. The same is valid for primary hair cells - similar to the mechanoreceptors present in neural animals -, which is quite enigmatic since Porifera has no neurons, although they present sensorineural genes (Mah and Leys 2017). Mah and Leys suggest that these markers may not be expressed in structures with sensory functions in Porifera, which would indicate a cooptation by eumetazoan nerve cells of sensory modules already present in non-neurological lineages.

Eumetazoa is here characterized by the presence of nervous system, neurotransmitters Glycine, Dopamine, Norepinephrine, Epinephrine and Serotonin (Karhunen et al. 1993; Liebeskind et al. 2017; Byrne 2019), endogenous opioid receptors (Bergamo et al. 1992; Im and Galko 2012; Sha et al. 2013; Mills et al. 2016; Scanes and Pierzchala-Koziec 2018), microvilli present in sensory cells (Manley and Ladher 2008) and support cells in mechanoreceptors (Fritzsch and Beisel 2001; Manley and Ladher 2008). The presence of neurotransmitters such as dopamine, which is a strong modulator of the motor system and acts on learning behaviours to rewards or punishments (Barron et al. 2010), indicates the existence of sensations through valences in Deuterostome, Protostome and non-bilateral animals with nerve cells. Our results show that even animals without centralization of the nervous system seem to experience and respond behaviourally to different stimuli through rewards and punishment.

The nociceptors and endogenous opioid receptors also point to a system of detection and responses to distinct harmful stimuli, probably originated in the common ancestor of Bilateria (Smith et al. 2009; Sneddon, 2017). The detection, avoidance and sometimes analgesia of harmful stimuli would be advantageous and, possibly, an adaptive characteristic. On the other hand, the opioid receptors known as ancient signaling systems (Im and Galko 2012; Mans et al. 2012; Guo et al. 2013) used by a multitude of living creatures generally seem to preserve their essential and protective reaction control functions (Raffa et al. 2013; Sha et al. 2013; Mills et al. 2016; Scanes and Pierzchala-Koziec 2018).

Our phylogenetic reconstruction of Bilateria supported by the presence of a ventrally organized nervous system with a longitudinal nerve cord, a cerebral ganglion, glial cells, amiellinic nerve fibers (Sneddon 2003; Ashley et al. 2007; Crook et al. 2013; Walters 2018; Wong and Rankin 2019) and horizontal commissures between the nerve cords, corroborate previous work (e.g. Smith and Lewin 2009; Sneddon 2017; Feinberg and Mallat 2016). The evidence of common ancestry reinforces the sharing of functions preserved in the acquisition, processing, and transport of information and stimuli in protostome and deuterostome nervous systems among Bilateria (Strausfeld and Hirth 2016; Ortega and Olivares-Bañuelos 2020). Even so, many events of secondary loss - of traits related to the nervous and sensory system may have happened during the evolution of some animal lineages such as Bivalvia, Echinodermata and Urochordata (Strausfeld and Hirth 2016).

According to our results, the nervous system is a synapomorphy of Eumetazoa, which is partially corroborating Philippe (2009)’s and Paulin and Cahill-Lane (2019)’s hypotheses. However, the view of a unique nervous system originating in the common ancestor of Eumetazoa is against recent genomic data that positioned Ctenophora as the sister-group of all the other Metazoa (Giribet 2016; Giribet and Edgecombe 2020), which would lead to at least two independent origin of nerve cells. The scenario proposed here supports previously reported assumptions, where the evolution of nervous systems has been considered to be codependent to other characteristics such as external digestion (Evans et al. 2019; Mángano and Buatois 2020) and motility (Paulin and Cahill-Lane 2019). These two characteristics together, according Paulin and Cahill-Lane (2019), created the possibility of meetings between different species in the environments, accompanied by the emergence of ecological relationships, such as predation. In this context some attributes may have been selected for more efficient escape from predation, and one of these characteristics that remains in almost all living animals, is the ability of flexible escape behaviours opportune by nervous systems. In this context, the conservation of biophysical mechanisms present in nervous systems of distinct species may be supported by different events in the early evolution of the nervous system (Paulin and Cahill-Lane 2019; Mángano and Buatois 2020).

The evolution of the centralization of the nervous system remains a controversial issue in evolutionary biology (Arendt et al. 2016). Martinez and Sprecher (2020) explain that the emergence of the nervous system through the neural condensation occurred by the presence of different types of sensory characteristics (e.g., photo and chemoreceptors), especially in the anterior part of the bilateral animals body, which would enable a more refined direction in the locomotion and an almost immediate perception of a new environment or stimulus (Martinez and Sprecher 2020). Our results point that Urbilateria had a centralized nervous system - according to Arendt et al. (2016), Cnidaria has nervous networks that, although diffuse, present genetic optimizations for centralization and specific functions. If such attribute was already present in the common ancestor of Eumetazoa, it is possible to explain the evolution of the centralized nervous system from a diffuse nervous network. Other evidence supporting such a scenario was found by Strausfeld and Hirth (2013) who observed similarities suggesting deep homology between the Arthropoda brain ganglia and the Vertebrata basal ganglia (Strausfeld and Hirth 2013; Held 2017). In this overview, the absence of centralization of the nervous system in some lineages of Bilateria may be due to secondary losses (Held 2017).

### 4.2. Behaviours related to sentience in Metazoa

The ability to detect harmful chemical, mechanical and thermal stimuli is essential for an adaptive life (Ginsburg and Jablonka 2010). Here, we suggest behavioural attributes are ubiquitous in animals. Associative learning allows connections between certain behaviours and their consequences, and also provides adaptive advantages related to pattern recognition and decision-making (Van Duijn 2017). Based on our results, classical conditioning is widely distributed among the Cnidaria, Ecdysozoa, Spiralia and Deuterostomia. On the other hand, operant conditioning is more restrictively distributed among Mollusca and Arthropoda, but widespread in Vertebrata, partially corroborating Cabanac et al. (2009)’s hypothesis. These results support two separate views: (1) that the different forms of detection of harmful stimuli and possibilities for associative learning were subject of different selection processes during the evolution of metazoan lineages (Van Duijn 2017), or (2) that detection and response behaviours derived from a common animal ancestor equipped with sensory-motor machinery which was able to actively respond to stimuli (Strausfeld and Hirth 2016).

The attribution of homologies for different behaviours in animals, however, should be carefully analyzed (Wenzel 1992; Van Duijn 2017). Further studies are necessary to understand the evolutionary patterns and processes related to the emergence and differentiation of behavioural attributes, which are extremely unstable (Gittleman and Decker, 1994; Gutfreund, 2018).

Considering the phylogenetic hypotheses here discussed, it is possible to infer that the first neural systems - with nerve cells and sensory organs accumulating in cephalic regions - date back to bilateral organisms living in the transition from the Ediacaran and Cambrian periods. Ediacaran fossils indicate traces of burrows, suggesting the presence of avoidance behaviours, i.e., a certain degree of perception of stimuli (Strausfeld and Hirth 2016; Coutts et al. 2018). Based on the scenarios presented here, the capacity for associative learning may have been one of the driving forces behind the animal diversification in the Cambrian as well as the emergency of capacity for subjective experiences, i.e., sentience, that was already underway in evolution of animals with nervous systems (Ginsburg and Jablonka 2010; Strausfeld and Hirth 2016).

### 4.3. Conservation of molecular sequences related to nociception and pain

With the emergence of molecular, genomic and developmental studies, homology assignment has expanded, applying not only to morphological characters but also to genes and ontogeny (Strausfeld and Hirth 2016). In this context, our molecular phylogenies point to a broad distribution of the genes analyzed here, which are present in several lineages of Metazoa in different degrees of plasticity, often acting in a polymodal way on pain and nociception mechanisms (Hoffstaetter et al. 2018).

Deg Enac seems to present a function conserved in the mechanical nociception of Mammalia (Bem-Shahar 2011; Han 2012), Hexapoda (Himmel et al. 2017) and Nematoda (Han 2012). The same is valid for NomP, present in the aforementioned groups and also in Actinopterygii (Vriens et al. 2004; Shin et al. 2005), and Piezo, which was also found in Porifera (Table 2). Prdm12 seems to be fundamental to support of nociceptor neurons (Vervoort et al. 2015; Walters 2018), and acts on mechanical nociception in Hexapoda (Nagy et al. 2015; Walters 2018), while Pkd2 is associated with thermal nociception with cold (Ye et al. 2008; Turner et al. 2016). The TRP family is involved in broad nociception functions, both thermal (Dhaka et al. 2007; Kadowaki 2015; Peng et al. 2015a,b; Himmel et al. 2020), mechanical (Vennekens et al. 2012; Kadowaki 2015; Sun et al. 2020) and chemical (Gallio et al. 2011; Venkatachalam et al. 2014; Kadowaki 2015).

Based on these nociception markers, our results reinforce the existence of a gene elements toolkit(as concepted by Walters 2018) conserved among animals and involved in nociceptive responses and pain mechanisms. As previously reported by Toth and Robinson (2007), the presence of such toolkit can help us to understand the evolution of different behaviours. Our evidence also shows the possibility of cooptation of genes such as Piezo, which were already present in ancient non-neural animals.

In view of the vertebrate-centric bias present in current research related to sentience (Andrade and Santos 2019), it is not surprising that our analyses based on molecular characters have generated inconclusive phylogenetic positioning for some lineages. The absence of wider taxonomic sampling in evolutionary studies may prevent further advances in the understanding of comparative biology and evolution of characteristics related to the nervous system and associated characteristics (Dos Santos et al. 2020), as well as hamper biological conservation efforts.

## 5. Conclusion

Although remaining an unsolved puzzle, the phylogenetic studies carried out in the present paper add supporting evidence for the Last Universal Common Sentient Ancestor (LUCSA) hypothesis. Sentience is a phenomenon that may have emerged through genetic bricolage and cooptation processes, which fundamental blocks existing even before the appearance of nervous systems were rearranged and acquired new functions. In this sense, the sentience may be widely distributed in Eumetazoa and seems to be especially related to origin of Urbilateria at least 542 million years ago.

Here, we discussed evidence for the possibility of common genotypic and phenotypic ancestry of attributes related to sentience in animal groups. Future directions should explore aspects of life history and selective pressures for each clade, compare cognitive skills among the different lineages of Metazoa, and focus on non-model species and neglected groups of Metazoa. The present research opens paths for new transdisciplinary dialogues as a form to obtain robust knowledge on the evolution of animal sentience, increasingly filling gaps about our knowledge of the problem of other minds.

## Acknowledgments

The authors would like to thank Maria Carolina Manzano and Stephanie Sampronha for reading and suggestions. This study was financed by the CAPES (Coordenação de Aperfeiçoamento de Pessoal de Nível Superior, Brazil - Finance Code 001), CNPq (307662/2019-5) and FAPESP (2017/11768-8).

## Availability of data and material

The datasets are available in the following repository *https://github.com/michaellaandrade/database_evolutionsentience*

